# *Caenorhabditis elegans* as a Model to Dissect Pharmacokinetic and Pharmacodynamic Relationships of Gabapentinoids

**DOI:** 10.64898/2026.08.26.747285

**Authors:** Jabin Sultana, Jesus D. Castaño, Jérôme R. E. del Castillo, Francis Beaudry

**Author notes:** Corresponding author: Francis Beaudry, Ph.D., Professor of Analytical Pharmacology, Canada Research Chair in metrology of bioactive molecules and target discovery Département de Biomédecine Vétérinaire, Faculté de Médecine Vétérinaire Université de Montréal, 3200 Sicotte, Saint-Hyacinthe, QC Canada J2S 2M2.

## Abstract

Gabapentin (GBP) and pregabalin (PGB) are widely used gabapentinoids. Previously, we have demonstrated, for the first time, that GBP and PGB modulate the nociceptive response to noxious heat in *C. elegans* at an optimal concentration. In the current study, we use *C. elegans* and paired thermal nociception assays with direct internal drug concentration measurements to characterize the pharmacokinetic (PK)/pharmacodynamic (PD) relationship of both compounds. Neither drug altered baseline mobility or quadrant preference, confirming that behavioral effects reflected genuine antinociceptive action. Both GBP and PGB produced dose- and time-dependent reductions in thermal avoidance, with 500 µM exposures generating a biphasic, V-shaped time course in which suppression of thermal sensitivity deepened before partially reversing. This partial reversal occurred later with PGB than with GBP. Internal concentrations confirmed dose-dependent absorption and retention for both drugs, yet at 500 µM, internal drug levels remained elevated through 360 min even as behavioral avoidance recovered, indicating that the recovery limb reflects active counter-regulation rather than passive clearance, consistent with previously reported transcriptional and proteomic signatures. Exposure–response profiles were notably flat, suggesting a saturable pharmacodynamic ceiling. Molecular modeling revealed conserved electronic pharmacophores supporting shared α2δ engagement, alongside shape-descriptor differences that may contribute to divergent absorption kinetics. These findings position *C. elegans* as a valuable model for dissecting gabapentinoid PK/PD relationships. Beyond mechanistic insight, these findings support the continued investigation of *C. elegans* as a screening platform whose validation could help address the 3R (Replacement, Reduction, Refinement) principles guiding animal research.

## Introduction

Gabapentin (GBP) and pregabalin (PGB) are widely used gabapentinoid medications that were originally developed for epilepsy but are now prescribed mainly for neuropathic pain, with additional established use as adjunctive treatments for focal or partial-onset seizures (Calandre et al., 2016; Pottegård, Anton et al., 2025; Greenblatt & Greenblatt, 2018; Taylor et al., 2007). Moreover, PGB is also used for generalized anxiety disorder in some clinical contexts (Baldwin et al., 2011; Fagan & Baldwin, 2023). Although gabapentin and pregabalin are structural analogs of GABA, their therapeutic effects do not appear to be mediated by GABA receptors (Taylor et al., 2007). Instead, experimental studies indicate that gabapentin, and other gabapentinoids, primarily targets the α2δ subunit of voltage-gated calcium channels, leading to reduced calcium influx and decreased release of excitatory neurotransmitters such as glutamate and aspartate (Field et al., 2006; Gee et al., 1996; Fink et al., 2002).

GBP and PGB are structurally related synthetic analogues of GABA that share a similar pharmacological target but they are not pharmcokinetically interchangeable (Ben-Menachem, et al., 2004; Bockbrader, Radulovic, et al., 2010; Bockbrader, Wesche, et al., 2010). GBP undergoes saturable intestinal absorption and displays nonlinear pharmacokinetics (Gidal et al., 1998; Stewart et al., 1993), while PGB behaves differently: it is absorbed more rapidly, shows high oral bioavailability, and tends to produce more predictable, dose-proportional exposure across the therapeutic range (Bockbrader, Wesche, et al., 2010; Lal et al., 2021). These differences provide an informative comparative framework. GBP and PGB share a common primary target, the α2δ-1 auxiliary subunit of voltage-gated calcium channels, but they are not pharmacodynamically equivalent: pregabalin binds α2δ-1 with higher affinity and additionally engages the α2δ-2 subunit, an interaction linked to altered cerebellar function. Divergent pharmacodynamic outcomes therefore reflect both target-level properties and differences in absorption, distribution, and elimination (Bockbrader, Wesche, et al., 2010; Mayoral et al., 2025; Zhou et al., 2024). In translational pharmacology, drug response is shaped not simply by the administered dose but by the concentration that actually reaches the biological system and by how that concentration changes over time (Bockbrader, Wesche, et al., 2010). Correlating internal drug concentration with pharmacodynamic effects is therefore a foundational principle of PK/PD analysis (Bockbrader, Wesche, et al., 2010). This relationship has been studied extensively in mammalian systems but remains less well characterized in alternative whole-organism models, such as those used for high-throughput screening (HTS). In these models, nominal exposure concentrations may not accurately reflect the actual internal concentration or the bioavailable fraction responsible for producing biological responses (Burns et al., 2010; Hartman et al., 2021; Zheng et al., 2013). Under such conditions, behavioral outcomes alone can be difficult to interpret without some measure of internal exposure (van der Most et al., 2024).

Among alternative in vivo systems, *Caenorhabditis elegan*s (*C. elegans*) is an excellent candidate for whole organism-based high-throughput screening (HTS) and toxicological research (Giunti et al., 2021; Kaletta & Hengartner, 2006). The nematode is easy to maintain, has a short life cycle of approximately three days at 20°C, and can be studied at a scale that would be difficult to achieve in vertebrate models (Leung et al., 2008). Moreover, its nervous system is fully mapped at single-cell resolution (White et al., 1986), and many signaling pathways relevant to neurotransmission, ion-channel function, and xenobiotic handling are sufficiently conserved to support pharmacological investigation (Hartman et al., 2021; Leung et al., 2008). In addition, *C. elegans* exhibits quantifiable thermo-nociceptive behaviors that have been used for decades as a behavioral assay of nocifensive responses (Glauser et al., 2011; Wittenburg & Baumeister, 1999).

Mammalian pain processing relies heavily on voltage-gated calcium channels (VGCCs), and *C. elegans* possesses direct orthologs of the VGCC subunits, namely UNC-2 (corresponding to the pore-forming principal α1 subunit) and UNC-36 (corresponding to the auxiliary α2δ subunit), which interact to regulate calcium channel localization and signaling (Caylor et al., 2013; Saheki & Bargmann, 2009). Together, these shared characteristics indicate that the *C. elegans* model can effectively reflect aspects of complex mammalian nociception, making it a valuable system for assessing both pain pathways and drug efficacy (Giunti et al., 2021).

Our previous work showed that GBP and PGB impair thermal avoidance behavior in *C. elegans* (Sultana et al., 2025); in addition, disrupted VGCC signaling directly alters *C. elegans* responses to noxious thermal stimuli, providing robust evidence for conserved calcium-dependent mechanisms in heat nociception (Iliff & Xu, 2020; Sultana et al., 2025). However, phenotypic responses alone are insufficient to characterize drug kinetics. The nominal concentration in the exposure matrix is rarely equal to the concentration in the internal fluids or at the biophase of the target receptors. Under steady-state conditions and linear pharmacokinetics, however, internal concentrations remain proportional to the external concentration, such that graded changes in exposure produce proportional changes at the target site. (Burns et al., 2010; Zheng et al., 2013). In *C. elegans* drug uptake may be influenced by cuticular permeability, transport processes, bacterial food interactions, and compound-specific disposition kinetics (Hartman et al., 2021; van der Most et al., 2024). As a result, identical nominal concentrations can yield distinct internal exposures, meaning phenotypic variations may reflect differences in drug absorption or accumulation rather than true target engagement (Burns et al., 2010; Zheng et al., 2013). Although some studies have explored drug accumulation and toxicokinetics in *C. elegans* (Burns et al., 2010; van der Most et al., 2024), there is still a significant gap in directly linking internal drug concentrations to behavioral pharmacodynamic outcomes - particularly for analgesic agents like gabapentinoids.

The present study was designed to address this gap by characterizing the pharmacokinetic and pharmacodynamic profiles of GBP and PGB in *C. elegans* using a time-matched exposure–washout paradigm. Nematodes were exposed to GBP or PGB for 1 h, washed to remove residual external compound, and then maintained under drug-free conditions. At defined post-exposure time points, internal drug concentrations were quantified in nematode extracts by HPLC-MS/MS, while thermal avoidance behavior was assessed on the same experimental schedule. This approach allowed internal drug burden and behavioral response to be examined in parallel over time, rather than as separate endpoints. By correlating post-exposure internal drug concentrations with a measurable nocifensive response, the study aimed to elucidate the influence of pharmacokinetics on behavioral outcomes and, more broadly, to evaluate the viability of *C. elegans* as a predictive, early-stage PK/PD framework for analgesic discovery.

## Materials and methods

### Chemicals and reagents

All chemicals and reagents were purchased from the Canadian division of Fisher Scientific (Ottawa, ON, Canada) or MilliporeSigma Canada Ltd. (Oakville, ON, Canada). Gabapentin (GBP), pregabalin (PGB), ^13^C-labeled gabapentin, and ^13^C-labeled pregabalin were purchased from MilliporeSigma Canada Ltd.

### C. elegans strains

The wild type N2 (Bristol) isolate of *C. elegans* was used as a reference strain. N2 (Bristol) was purchased from the Caenorhabditis Genetics Center (CGC), University of Minnesota (Minneapolis, MN, USA). *C. elegans* was maintained and handled under standard conditions as previously published (Margie et al., 2013). Nematodes were grown and kept on nematode growth medium (NGM) agar at 22 °C in a Thermo Scientific incubator (Fair Lawn, NJ, USA). Experiments were performed at room temperature (∼ 22 °C) unless otherwise mentioned.

### Pharmacological manipulation

GBP and PGB stock solutions (1 mM) were prepared using Type 1 ultrapure water and mixed thoroughly by vortexing. Working solutions of 100 µM and 500 µM were subsequently prepared by serial dilution of the stock solutions with Type 1 ultrapure water. Wild-type *C. elegans* were cultured for 72 h on nematode growth medium (NGM) plates (92 × 16 mm) seeded with *Escherichia coli* OP50 as a food source. Adult worms were then exposed to GBP or PGB solutions at the desired concentrations.

Approximately 10 mL of drug solution was added to each plate, producing a thin liquid layer of approximately 2–3 mm, allowing the worms to remain immersed during exposure.

For both pharmacokinetic and behavioral experiments, nematodes were exposed to GBP or PGB for 1 h. Following exposure, nematodes were washed three to four times with S basal buffer to remove residual external drug and then transferred to drug-free conditions. Thermal avoidance behavior and pharmacokinetic sampling followed the same exposure–washout and post-exposure time-course design. Worms were maintained in drug-free buffer and collected at defined time points of 0, 15, 30, 60, 90, 120, 180, 240, and 360 min after the exposure period. Prior to each behavioral assay or pharmacokinetic collection, worms were washed again with S basal buffer to further minimize residual external drug contamination.

### Thermal avoidance assay

Thermal avoidance assays were performed at the same post-exposure time points used for pharmacokinetic analysis. Following drug exposure and washing, worms were maintained in drug-free conditions until behavioral assessment. The thermal avoidance assay was carried out as previously described by (Margie et al., 2013). Briefly, experiments were conducted on 92 × 16 mm Petri dishes divided into four quadrants. In one setup, all four quadrants were maintained at control room temperature (22–25 °C) without thermal stimulation to serve as a negative control. In another setup, two opposite quadrants were assigned to the control temperature (22–25 °C), while the remaining two were allocated for thermal stimulation (32–35 °C). The plate layout comprised a central circle 1 cm in diameter, from which *C. elegans* was excluded, and four quadrants: two stimulation regions (A and D) and two control regions (B and C). Sodium azide (0.5 M) was applied to immobilize nematodes in all quadrants at a distance of 1.5 cm from the center, preventing their spread across the plate. An electronically heated metal tip (0.8 mm in diameter) generated noxious heat, producing a radial temperature gradient of 32– 35 °C on the NGM agar, measured 2 mm from the tip using an infrared thermometer. The stimulus temperature was based on a previous study (Sultana et al., 2025). Nematodes were collected and washed following the method of (Margie et al., 2013) after 72 h of growth on *E. coli*-seeded NGM plates. An aliquot of 2–3 μL (typically containing 100 to 500 young adult nematodes) was then placed at the center of a Petri dish within the 1 cm marked circle and subjected to thermal stimulation in the designated quadrants (A and D). After 30 min, the plates were removed and stored at 4 °C for a minimum of 1 h to immobilize all nematodes. The nematodes in each quadrant were then counted under a stereoscope, excluding those that had not crossed the 1 cm circle. Nocifensive responses to noxious heat were analyzed for each *C. elegans* group using the thermal avoidance index (TI) and thermal avoidance percentage, as detailed in Supplementary Figure S1.

### Sample Preparation for HPLC–MS/MS Analysis

For pharmacokinetic analysis, three independent biological replicates were prepared for each treatment condition and time point. For the T0 condition, nematodes were immediately homogenized after the initial washing step using ice-cold phosphate buffer to minimize any further metabolism or redistribution of the compounds. For subsequent time points, nematodes were maintained in fresh drug-free buffer until collection. Samples were homogenized in ice-cold phosphate buffer using glass beads and an electric homogenizer. Homogenates were centrifuged at 12,000 rpm for 10 min, and the resulting supernatants were collected for further analysis. Protein concentration in each sample was determined using a Bradford protein assay, and these values were later used to normalize measured drug concentrations using a protein-based correction factor. For HPLC–MS/MS analysis, 100 µL of each sample supernatant was mixed with acetonitrile containing the internal standard (i.e. ^13^C-labeled gabapentin (100nM), and ^13^C-labeled pregabalin (500nM)). Samples were vortex-mixed thoroughly and centrifuged at 12000 rpm for 10 mins to precipitate proteins. The resulting supernatants were collected and transferred into HPLC vials for HPLC–MS/MS analysis.

### Bioanalytical method

Samples (5 µL) were injected onto a Vanquish UHPLC system coupled to a TSQ Altis Plus triple-quadrupole mass spectrometer (Thermo Scientific, Waltham, MA, USA). Chromatographic separation was achieved on a Hypersil Gold column (50 × 2.1 mm) maintained at 30 °C. GBP was separated under isocratic conditions with a mobile phase of 80% solvent A (0.1% formic acid in water) and 20% solvent B (0.1% formic acid in acetonitrile) at a flow rate of 0.2 mL/min over a 2-min run. PGB was separated using a linear gradient of solvent A (0.1% formic acid in water) and solvent B (0.1% formic acid in acetonitrile) at a flow rate of 0.3 mL/min. The gradient was programmed as follows: 0 min, 5% B; 3.5 min, 95% B; 3.9 min, 95% B; followed by re-equilibration to 5% B.

Mass spectrometric detection was performed in positive ionization mode using multiple reaction monitoring (MRM). Source parameters were held constant across all analyses: spray voltage, 3500 V; sheath gas, 40 arb; auxiliary gas, 5 arb; sweep gas, 0.5 arb; ion transfer tube temperature, 325 °C; and vaporizer temperature, 350 °C. The collision-induced dissociation (CID) gas pressure was maintained at 2 mTorr. The collision energy was 20 V for all transitions. Additional acquisition parameters were: chromatographic peak width, 8; dwell time, 100 ms; calibrated RF lens enabled; Q1 resolution, 0.7 FWHM; and Q3 resolution, 0.7 FWHM. The following MRM transitions were monitored: gabapentin, precursor m/z 172 → product ions m/z 137 and 154; ^13^C_3_-gabapentin, precursor m/z 175 → product ions m/z 140 and 157; pregabalin, precursor m/z 160 → product ions m/z 125, and 142 and ^13^C_3_-pregabalin, precursor m/z 163 → product ions m/z 128, and 145. The quantification was based on specific MRM extracted ion chromatograms and analyte concentrations were determined using the peak area ratio of the light and heavy analogs.

### Preliminary Pharmacokinetic and pharmacodynamic analyses

PK parameters were determined for each experimental group using a one compartmental analysis (60 min infusion simulation) using PKSolver (Zhang et al., 2010). AUC of *in situ* concentration over time from time 0 to final timepoint (AUC_0–t_), AUC of concentration over time from time 0 to infinity (AUC_0–∞_), maximum concentration (C_max_), elimination half-life (T_1/2_) and mean residence time (MRT) was calculated.

### Molecular Modeling

Molecular modeling calculations were performed using Spartan’20 software (Wavefunction, Inc., Irvine, CA, USA) (Shao et al., 2006). Initially, three-dimensional space-filling (CPK) models of GBP and PGB were generated. A systematic conformational search was then conducted to identify the most stable conformer of each molecule, corresponding to the minimum-energy structure. Geometry optimization of the lowest-energy conformers was carried out using the Merck Molecular Force Field (MMFF) method (Halgren, 1996) and Hartree-Fock 6-31G*. Subsequently, the optimized geometries were used to calculate molecular properties and topological descriptors. Quantum chemical calculations were performed using Density Functional Theory (DFT) with the hybrid B3LYP functional (Brinzei et al., 2020) and the 6-31G* polarized basis set (Gill et al., 1992; Tirado-Rives & Jorgensen, 2008). All calculations were conducted in water and corresponded to the equilibrium geometry of the molecules in their electronic ground state.

### Statistical analysis

Behavioral data was analyzed utilizing the non-parametric Kruskal–Wallis followed by the post-hoc Dunn test for multiple comparisons. The significance level was set at p ≤ 0.05. Statistical analyses were conducted utilizing GraphPad PRISM (version 11.0.2).

## Results and discussion

We first examined whether the experimental setup itself affected worm behavior. Wild-type N2 worms were tested for mobility and quadrant preference, with or without exposure to gabapentin (GBP) or pregabalin (PGB). During this control experiment, all four quadrants were kept at room temperature, approximately 22 °C. As shown in Fig. 2, the worms did not show a preference for any particular quadrant, regardless of drug exposure or drug concentrations, 100 and 500 µM. After 30 minutes of being placed at the center of the Petri dish, the worms remained evenly distributed across the four quadrants. Exposure to GBP or PGB for 60 min also had no significant effect on their overall mobility.

**Figure 1.**
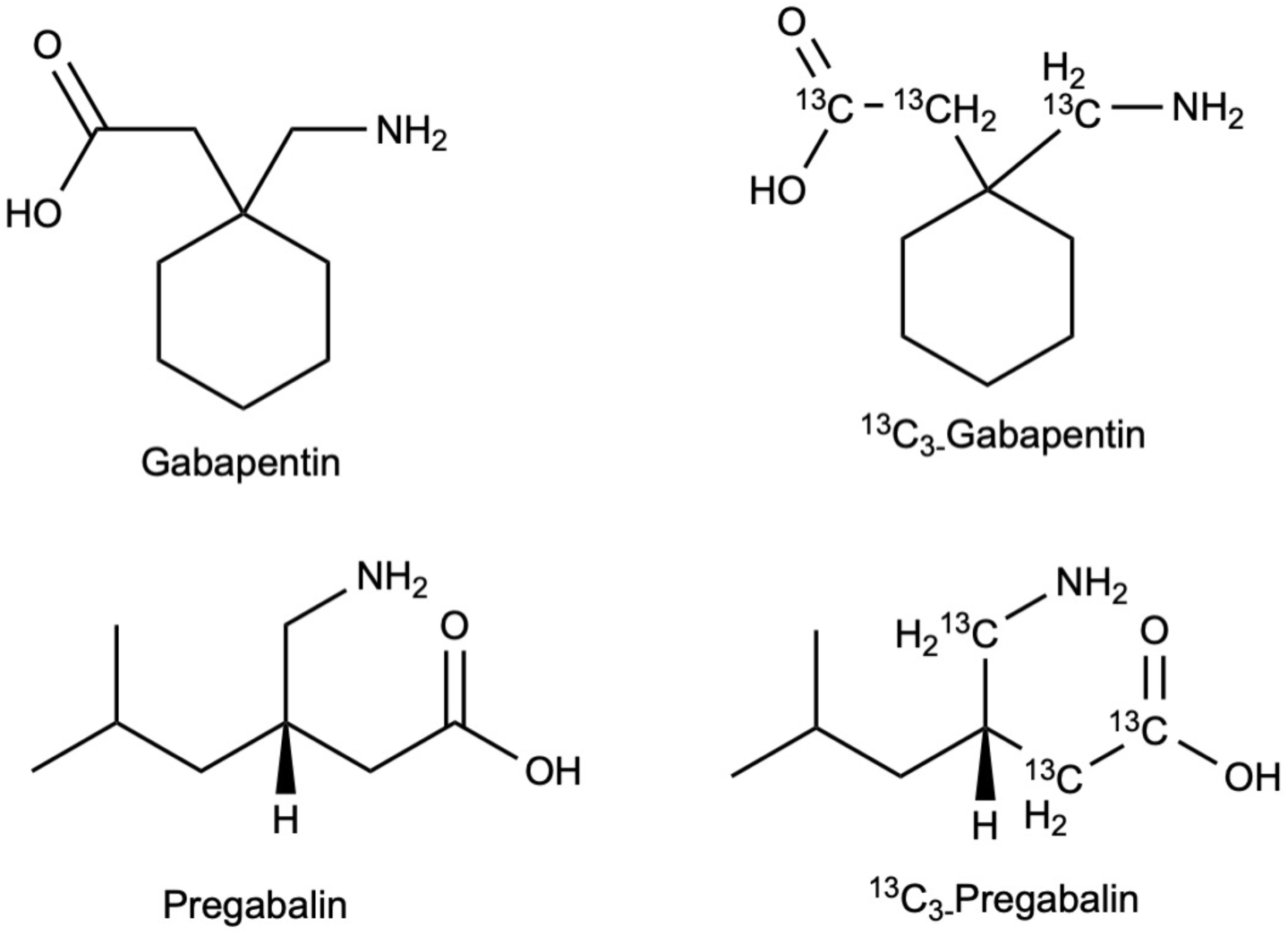
Structures of gabapentin, pregabalin, and their stable-isotope homologs, selective ligands of voltage-gated calcium channels containing the α2δ-1 subunit.

**Figure 2.**
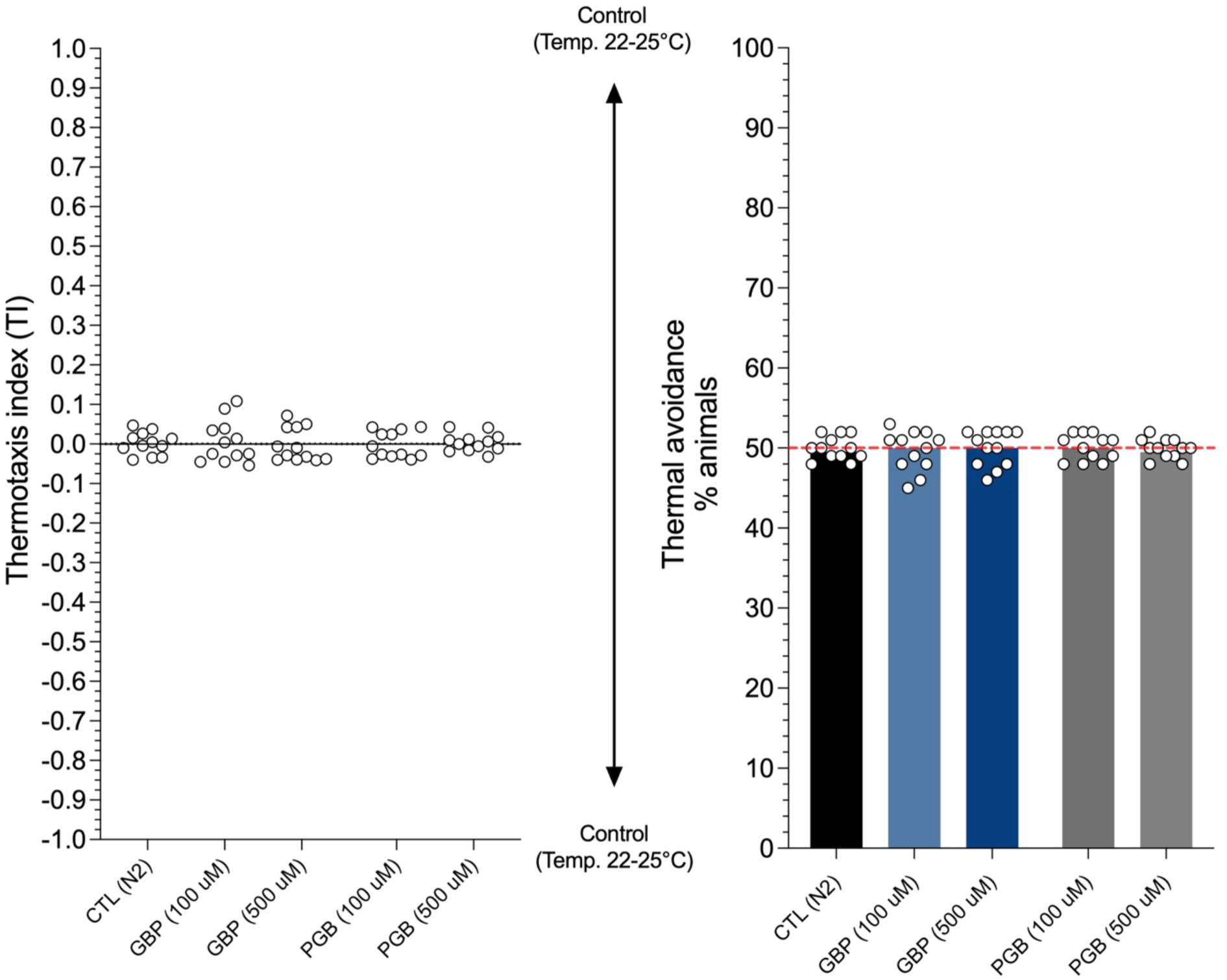
Comparison of the mobility and bias of WT (N2) and all selected mutant strains in petri dishes with NGM, divided into quadrants conserved at a room temperature (22 °C) without the application of a stimulus (negative control). No quadrant selection bias was observed for any *C. elegans* genotype tested in the absence or presence of PGB or GBP. Individual values and medians are displayed and derived from at least 12 independent experiments for each experimental group. (n = 100 to 500 nematodes per petri)

### Evaluation of Antinociceptive Activity of Gabapentin and Pregabalin

Both gabapentin and pregabalin produced a robust, time-dependent reduction in thermal avoidance behavior in *C. elegans*. In control worms, the thermotaxis index (TI) was strongly negative, reflecting pronounced avoidance of the noxious zone (32–35 °C). Following a 60-min exposure to 100 µM gabapentin, TI shifted markedly toward zero at all post-exposure time points (Fig. 3A), indicating an attenuated nocifensive response. This effect was maximal early (0–30 min; p < 0.0001) but partially diminished thereafter, yet remained significant through 360 min (p < 0.01). A parallel decline was observed in the proportion of heat-avoiding animals, from ≈78% in controls to ≈60–67% across treated groups (Fig. 3B). At 500 µM, gabapentin produced a biphasic, V-shaped time course (Fig. 3 C,D). The antinociceptive effect was modest immediately after washout and deepened to a maximum near 60 min. At peak effect, the thermotaxis index shifted significantly toward zero and the proportion of heat- avoiding animals reached its minimum, indicating a marked loss of heat avoidance. This response was only partially sustained, with avoidance behavior returning toward baseline by 360 min. The descending limb is consistent with continued internal accumulation and α2δ target engagement, whereas the ascending limb more likely reflects active counter-regulation than simple drug clearance. This interpretation links our present kinetic data with our earlier proteomic findings (Sultana et al., 2025), in which 500 µM gabapentin, unlike the 100 µM dose, upregulated translation, ribosomal biogenesis, and Wnt signaling: compensatory programs that favor pro-neuropeptide synthesis and neuronal hyperexcitability and could progressively oppose antinociception, producing the recovery limb of the V.

**Figure 3.**
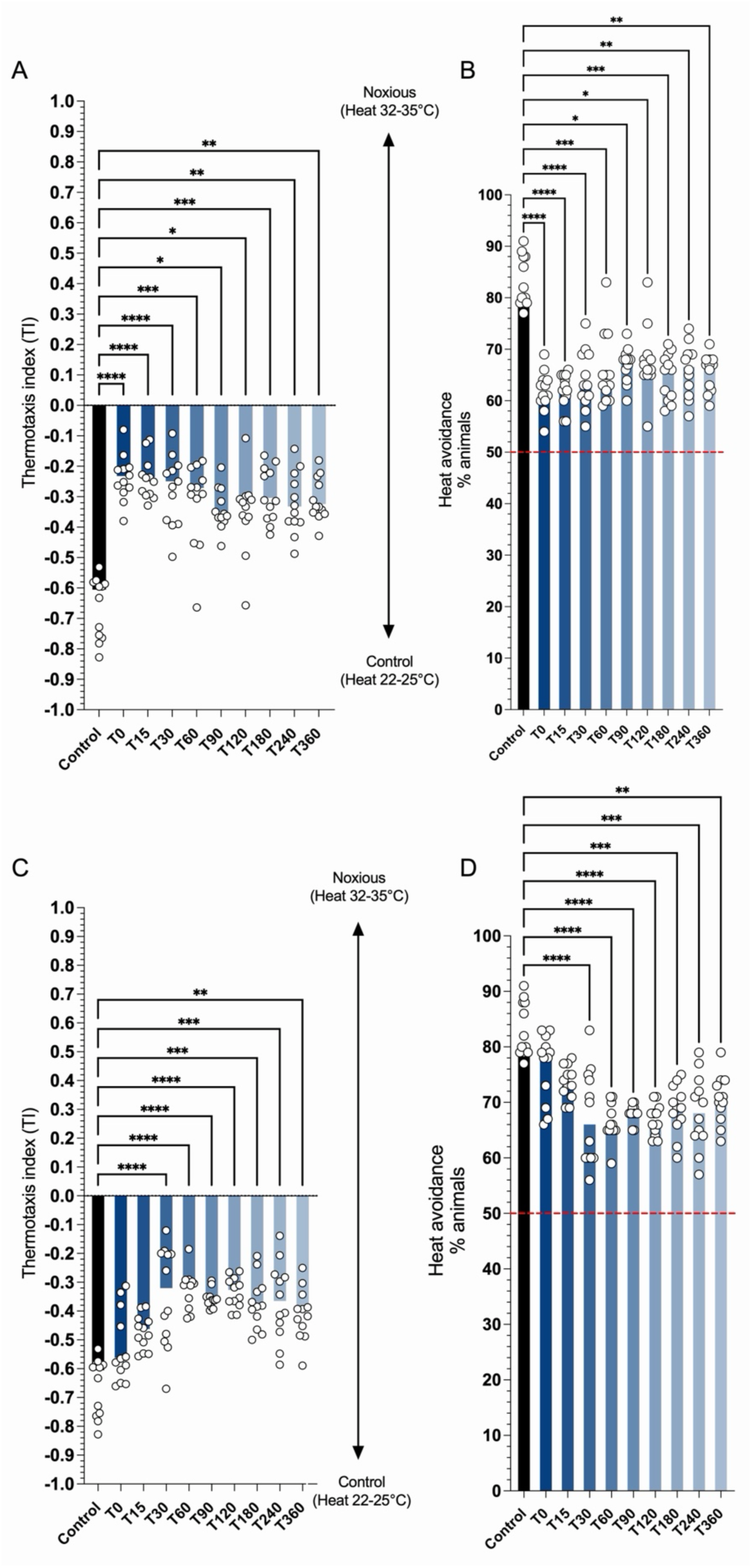
Assessment of gabapentin’s pharmacological impact on thermal avoidance behavior in *C. elegans.* Data represent individual values with medians from at least 12 independent experiments. Animals were incubated with 100 µM (A,B) or 500 µM (C,D) gabapentin for 60 min, then evaluated for time-dependent behavioral responses. Statistical analyses were performed using the Kruskal–Wallis test followed by Dunn’s multiple-comparison test (n = 100–500 nematodes per plate).

Comparing the two doses, both concentrations produced significant, time-dependent antinociception, but their kinetic profiles differed. At 100 µM the strongest effect was concentrated in the earliest post-exposure interval, whereas 500 µM produced a biphasic, V-shaped time course in which heat avoidance suppression deepened to a maximum near 60 min before partially reversing as avoidance behavior returned toward baseline. This graded, exposure-driven profile argues for a concentration- and time-dependent action rather than an all-or-none response. Critically, the recovery limb at 500 µM is unlikely to reflect simple drug clearance alone; it is more consistent with progressive counter-regulation, since the internal drug concentration remained substantial across the time course while avoidance recovered. This pattern underscores the value of pairing behavioral readouts with direct pharmacokinetic measurement and reinforces *C. elegans* as a tractable system for interrogating analgesic PK/PD relationships.

Pregabalin reproduced the overall pattern seen with gabapentin, with dose-specific kinetic differences. In control animals the thermotaxis index was strongly negative with most of nematodes avoiding the noxious zone. At 100 µM, PGB attenuated avoidance at all post-exposure time points (Fig. 4 A,B), shifting TI toward zero and reducing the heat-avoiding fraction significantly. As with GBP, the effect was maximal early (0–15 min; p < 0.0001) and gradually weakened over the time course, with significance decline to p < 0.05 by 240–360 min. At 500 µM, PGB again produced a biphasic, V-shaped profile (Fig. 4 C,D): heat avoidance suppression deepened to a low level before partially reversing as avoidance returned toward baseline by 360 min. Notably, the reached suppression level appeared later than with GBP, around 90–120 min rather than 60 min, and the recovery limb was correspondingly shifted.

**Figure 4.**
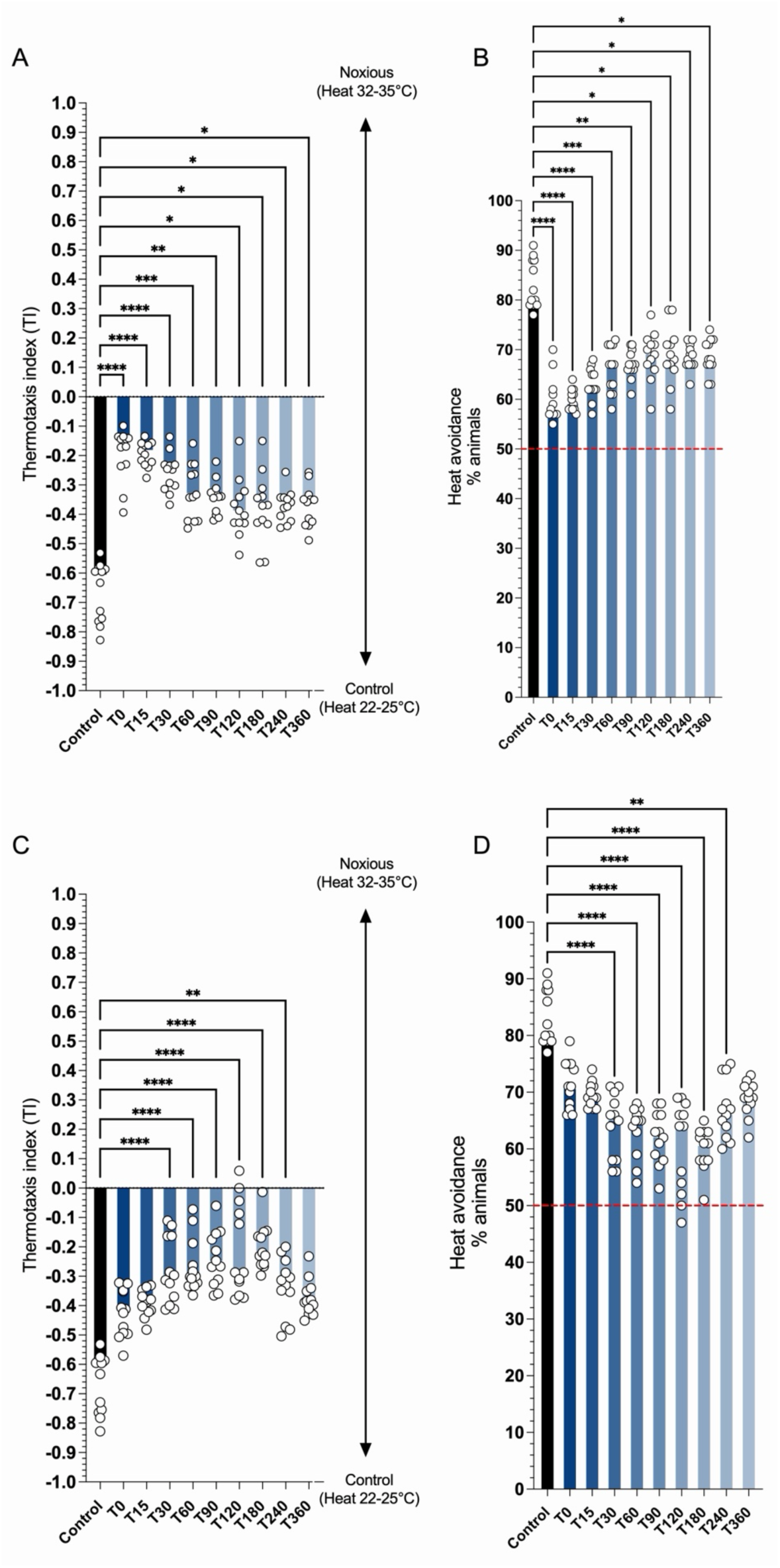
Pharmacological effects of pregabalin on thermal avoidance in *C. elegans*. Individual data points and medians are shown, pooled from ≥12 independent experiments per condition. Nematodes were exposed to 100 µM (A,B) or 500 µM (C,D) pregabalin for 60 min, followed by behavioral testing at multiple time points. Statistical significance was assessed using the Kruskal–Wallis test with Dunn’s post-hoc correction (n = 100–500 nematodes per plate).

The similarities are striking, both gabapentinoids showed early-maximal effects at 100 µM and a biphasic high-dose response, supporting a shared, exposure-driven mode of action consistent with α2δ engagement. The main difference was temporal, PGB’s high-dose maximal antinociceptive effect was delayed relative to GBP, which may reflect the distinct absorption and distribution kinetics of the two compounds. This mirrors their divergent pharmacokinetics in mammals and reinforces the notion that a shared mechanism can yield different behavioral time courses through PK rather than PD differences.

### Preliminary Pharmacokinetics and gabapentinoids heat avoidance effect

Following a 60 min drug exposition, internal drug concentrations confirmed that both gabapentinoids were absorbed and retained in *C. elegans* in a dose-dependent manner (Fig. 5 A,C). As shown in Table 1, both GBP and PGB exhibited dose-dependent increases in internal concentration, with C_max_ rising approximately 7- to 8-fold between the 100 µM and 500 µM exposures for each drug. GBP displayed relatively stable elimination kinetics across concentrations, with T_½_ (247.7 vs. 263.8 min) and MRT (357.3 vs. 380.6 min) remaining similar at both doses, suggesting first-order elimination within this concentration range. Pregabalin, in contrast, showed a marked prolongation of T_½_ (721.4 min at 100 µM vs. 18,398.1 min at 500 µM) and MRT. This divergence between observed and extrapolated exposure at 500 µM PGB suggests the terminal elimination phase may be poorly defined by the sampling window, and the resulting T_½_, MRT and AUC_0-∞_ values should be interpreted cautiously pending confirmation of the terminal-phase regression fit. Important to note, Terminal half-life (T_½_) and mean residence time (MRT) could not be adequately determined with the current model. Sampling beyond 6 h was precluded by the nematode replication rate, which would progressively alter the biomass of the exposed population and thereby confound the interpretation of measured internal concentrations. Because the terminal phase was consequently not characterized over a sufficient number of half-lives, λ_z_-dependent parameters, including T_½_ and MRT, should be regarded with caution. Alternative modeling approaches accounting for this sampling constraint are currently being investigated.

**Figure 5.**
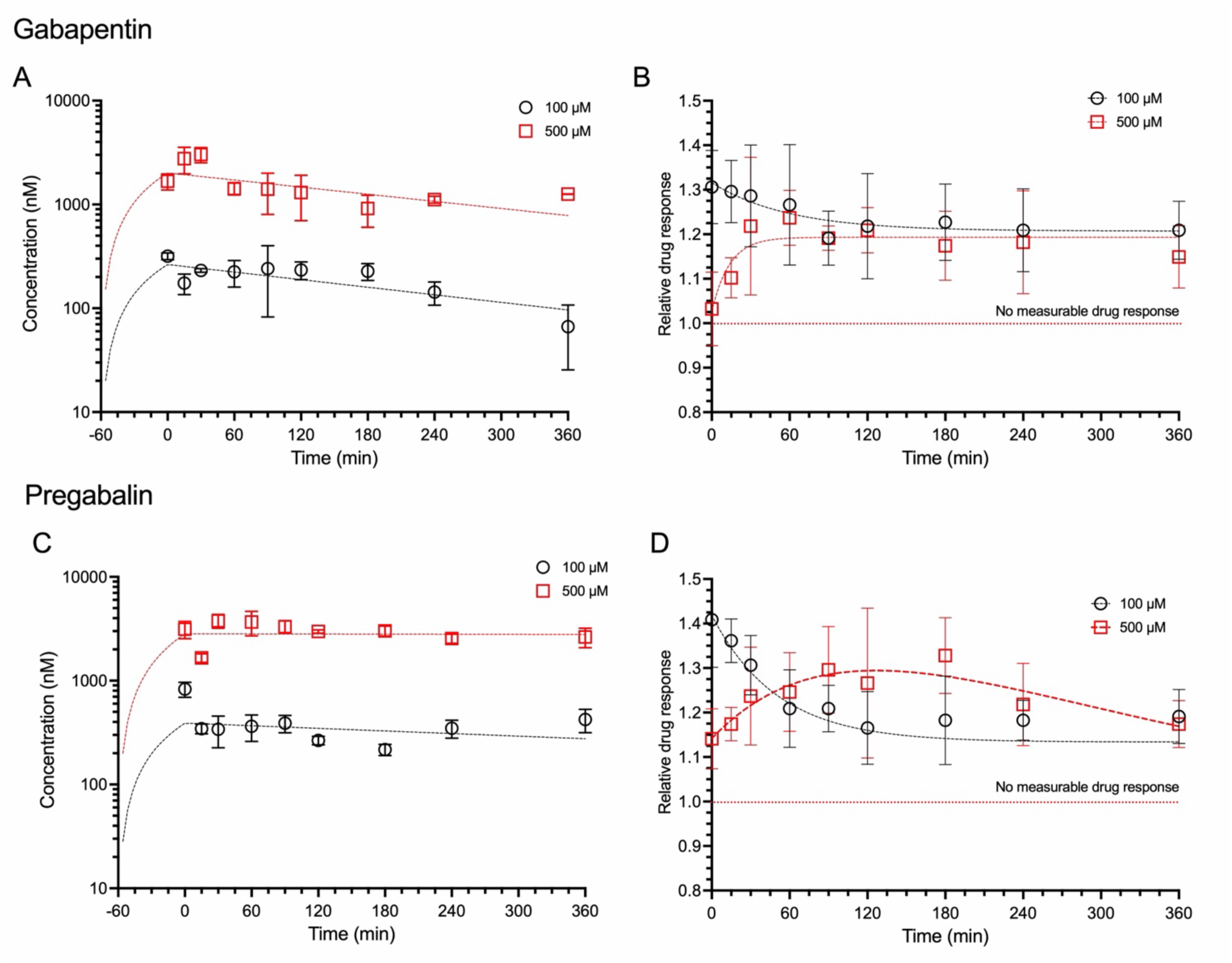
Concentration–time profiles and behavioral response kinetics of gabapentin (A,B) and pregabalin (C,D) in *C. elegans*.

**Table 1.** Preliminary pharmacokinetic parameters of gabapentin and pregabalin in *C. elegans* following 60-minute exposure in solution.

| Pharmacokinetic parameter | GBP 100μM | GBP 500 μM | PGB 100 μM | PGB 500 μM |
| --- | --- | --- | --- | --- |
| C<sub>max</sub> (nM) | 263.9 | 2011.6 | 389.9 | 2825.2 |
| T<sub>1/2</sub> (min) | 247.7 | 263.8 | 721.4 | 18398.1* |
| AUC<sub>0-t</sub> (nM *min) | 68010.4 | 530219.1 | 130472.3 | 1094981.1* |
| AUC<sub>0-∞</sub> (nM *min) | 102442.1 | 827479.6 | 417595.0 | 75072886.8* |
| MRT (min) | 357.3 | 380.6 | 1040.7 | 26542.9* |

Note: T<sub>1/2</sub>, AUC<sub>0- ∞</sub> and MRT could not be reliably estimated under the present design; refinement of the kinetic model to address this limitation is currently under investigation.

At 500 µM, internal concentrations of both GBP and PGB rose rapidly and remained elevated and essentially stable across the full 360-min window, whereas the 100 µM exposures produced correspondingly lower internal concentration and a somehow expected decline with time, especially for GBP. Critically, at 500 µM the internal concentration did not decline appreciably by 360 min, yet the behavioral response (Fig. 5 B,D) recovered toward baseline over the same interval. This temporal divergence, sustained internal drug alongside a diminishing behavioral effect, is the key finding: the recovery limb of the high-dose V-shaped response cannot be explained by simple pharmacokinetic clearance and instead points to active counter-regulation, consistent with the translation, ribosomal- biogenesis, and Wnt/SUMOylation related processes we previously identified at 500 µM (Sultana et al., 2025). The relative-response profiles also expose a striking dissociation between exposure and effect. At both doses and for both drugs, relative responses clustered around similar values despite approximately ten-fold differences in measured internal concentration. This dissociation is expected rather than anomalous. Whole-organism homogenate concentrations do not necessarily reflect concentrations at the target site, and the concentration–effect relationship is nonlinear. As concentrations increase beyond the EC50 and approach saturation, further increases in drug exposure produce progressively smaller changes in response. This somehow flat exposure–response relationship may indicate a saturable, ceiling-limited mode of action at the target rather than a graded one. Notably, at 500 µM both compounds showed a delayed rise to maximal response before declining, again decoupling the effect from the stable internal concentration and reinforcing that pharmacodynamic outcome, not exposure alone, governs the behavioral time course. Both features of our exposure–response data have precedent in mammalian pharmacology. For gabapentin, dose escalation does not reliably produce proportional gains in analgesic effect, and this pattern is commonly attributed, at least in part, to saturable intestinal absorption and declining oral bioavailability as dose increases (Cundy et al., 2008; Fabritius et al., 2017). In humans, gabapentin bioavailability decreases from roughly 60% to 33% across the therapeutic dose range, and plasma exposure rises less than proportionally with dose (Bockbrader, Wesche, et al., 2010; Cundy et al., 2008; Stewart et al., 1993). Because internal concentrations were measured directly here, and the response remained flat across an approximately ten-fold concentration range, the observed ceiling is more consistent with a downstream pharmacodynamic limit than with restricted uptake alone. However, it is not possible to establish that biophase concentrations scaled proportionally, since whole-organism homogenate represents an aggregate across tissues and cannot resolve the concentration at the target receptors. A delayed rise to maximal effect is likewise consistent with prior pregabalin PK/PD studies in rats: brain microdialysis experiments showed a counter-clockwise hysteresis between brain extracellular concentration and anticonvulsant effect, with response lagging behind measured drug levels and not being directly proportional to extracellular concentration (Feng et al., 2001). That interpretation also fits the broader pharmacology of pregabalin, which otherwise shows rapid, linear, and predictable systemic pharmacokinetics, indicating that delayed effect can persist even when exposure is well behaved (Darlami & Sharma, 2024). Together, these parallels suggest that the concentration–effect decoupling observed in *C. elegans* reflects a conserved feature of gabapentinoid pharmacodynamics rather than a purely species-specific artifact.

### Molecular modeling

Further structural and physicochemical analyses were performed as discussed in (Darlami & Sharma, 2024; Nkambeu et al., 2021). Key parameters for quantitative structure-property relationships (QSPR) and quantitative structure-activity relationships (QSAR) analysis are reported in Table 2. The computed molecular descriptors revealed that gabapentin and pregabalin are electronically near-equivalent at their shared pharmacophore. HOMO energies (-6.36 vs. -6.42 eV), dipole moments (2.08 D, identical), and polarizabilities (54.98 vs. 54.56 ×10⁻³⁰ m³) showed minimal divergence, while HBD (1) and HBA (2) counts were conserved between both molecules. This supports a common electronic basis for α2δ subunit engagement, with the amine donor and carboxylate/carbonyl acceptors presenting equivalent H-bonding capacity to the binding pocket in both compounds (Meneses et al., 2021; Mondal et al., 2024; Taylor et al., 2007). The principal distinction emerged in the LUMO energy (0.10 vs. 0.25 eV), which propagated to a modestly wider HOMO–LUMO gap for pregabalin (6.67 vs. 6.46 eV). Given that this difference (∼3%) approaches the uncertainty of the B3LYP model, both compounds are best described as electronically comparable, consistent with the low metabolic lability shared by the two drugs, although that property is governed primarily by their poor recognition as CYP450 substrates rather than by frontier-orbital energetics (Ben-Menachem, 2004). Shape descriptors further differentiated the two scaffolds: pregabalin displayed higher ovality (1.37 vs. 1.30) and surface area (212.30 vs. 205.84 Å²) despite a smaller molecular volume (182.01 vs. 186.52 Å³), reflecting its extended branched-chain conformation relative to gabapentin’s more compact, cyclohexane-constrained geometry. Correspondingly, gabapentin’s modestly higher LogP (0.88 vs. 0.75) is attributable to the additional lipophilic ring surface. These physicochemical differences may contribute to distinct steric fit within the α2δ amino-acid binding pocket, but in the absence of docking or binding-energy calculations, this should be regarded as a structural hypothesis rather than an established mechanism. Importantly, the well-documented absorption differences between the two drugs, saturable versus linear kinetics, are governed primarily by LAT1 transporter affinity (Belliotti et al., 2005; Bockbrader, Wesche, et al., 2010). These physicochemical differences may possibly extend beyond passive diffusion to influence carrier-mediated transport itself, as LAT1 substrate recognition is known to depend on side-chain bulk, shape, and lipophilicity; however, direct transporter kinetic data would be required to confirm a causal link to the observed saturable versus linear absorption kinetics of gabapentin and pregabalin, respectively. More specifically, this divergence may reflect differences in both transporter affinity indirectly measured with the Michaelis-Menten constant, K_m_ and maximal transport velocity (V_max_) between the two compounds.

**Table 2.** Predicted molecular properties for gabapentin and pregabalin, using DFT method, B3LYP model using the 6-31G* basis set, in vacuum, for equilibrium geometry at ground state.

|  | Gabapenti<br>n | Pregabalin |
| --- | --- | --- |
| **Molecular properties** |  |  |
| Formula | C<sub>9</sub>H<sub>17</sub>NO<sub>2</sub> | C<sub>8</sub>H<sub>17</sub>NO<sub>2</sub> |
| Molecular weight (amu) | 171.241 | 159.230 |
| Energy (au) | -558.425103 | -520.317093 |
| Dipole moment (debye) | 2.08 | 2.08 |
| E HOMO (eV) | -6.36 | -6.42 |
| E LUMO (eV) | 0.10 | 0.25 |
| $\Delta E$ (\| E<sub>HOMO</sub>–E<sub>LUMO</sub> \|) (eV) | 6.46 | 6.67 |
| **QSAR properties from CPK model** |  |  |
| Area (Å<sup>2</sup>) | 205.84 | 212.30 |
| Volume (Å<sup>3</sup>) | 186.52 | 182.01 |
| PSA (Å<sup>2</sup>) | 57.238 | 59.335 |
| Ovality | 1.30 | 1.37 |
| **QSAR properties from computed electron density** |  |  |
| Log P | 0.88 | 0.75 |
| HBD count | 1 | 1 |
| HBA count | 2 | 2 |
| Polarizability (10<sup>-30</sup> m<sup>3</sup>) | 54.98 | 54.56 |

HOMO and LUMO isosurfaces provide an approximate map of electron-rich (nucleophilic) and electron-deficient (electrophilic) regions within the molecule (Plasser & González, 2016; Yoğurtçu & Ersanlı, 2025). In gabapentinoids, this partitioning places the amine within the principal donor region and the carboxylate/carbonyl group within the principal acceptor region, supporting plausible electrostatic and hydrogen-bonding interactions with the α2δ binding site, although frontier orbital topology alone does not determine binding affinity or binding energy (Meneses et al., 2021). The HOMO-LUMO gap is commonly interpreted as an index of electronic stability and chemical reactivity, and therefore may serve as an indirect descriptor of metabolic resilience rather than a direct predictor of pharmacological potency (Bendjeddou et al., 2016). Likewise, polarizability and overall charge distribution are relevant physicochemical descriptors that can influence lipophilicity, transporter recognition, and renal handling, but these relationships remain indirect and should not be interpreted as single-cause determinants of pharmacokinetic behavior (Bendjeddou et al., 2016).

DFT-derived frontier molecular orbitals (i.e. B3LYP/6-31G*) shown in Figure 6 revealed a conserved spatial partitioning of electron density in both gabapentinoids: the HOMO localized predominantly over the primary amine nitrogen and adjacent alkyl framework, while the LUMO localized over the carboxylic acid/carbonyl moiety, with negligible overlap between the two isosurfaces in either molecule. This pattern is consistent with the shared zwitterionic pharmacophore required for α2δ subunit recognition, in which the protonated amine and deprotonated carboxylate engage the binding pocket as spatially independent electrostatic termini. The conserved HOMO/LUMO topology between gabapentin and pregabalin therefore supports, at the orbital level, the structural rationale for their common mechanism of action, rather than indicating a difference in receptor affinity (Meneses et al., 2021; Sinha et al., 2013). While the HOMO isosurfaces of gabapentin and pregabalin display an apparent inversion in phase (color) assignment, this reflects the arbitrary sign convention inherent to independently computed wavefunctions rather than a physically meaningful electronic difference; the relevant nodal topology (Takatsuka & Arasaki, 2021). Scaffold-dependent differences were nonetheless apparent: pregabalin’s HOMO lobe over the amine appeared more diffuse and extended toward the α-carbon, plausibly reflecting reduced steric constraint from its branched isobutyl chain relative to gabapentin’s cyclohexane ring. Whether this translates into a functionally relevant difference in electron availability at the amine, and by extension into altered receptor engagement, cannot be established from orbital topology alone and would require docking or binding-energy calculations to substantiate. Critically, the established differences in absorption between the two drugs, saturable, transporter-limited uptake for gabapentin via LAT1 versus linear, dose-proportional absorption for pregabalin, are governed by whole-molecule fit within the transporter binding site (charge spacing, steric bulk) rather than by frontier orbital energetics, and should not be attributed causally to the HOMO/LUMO differences observed here (Hutchinson et al., 2022).

**Figure 6.**
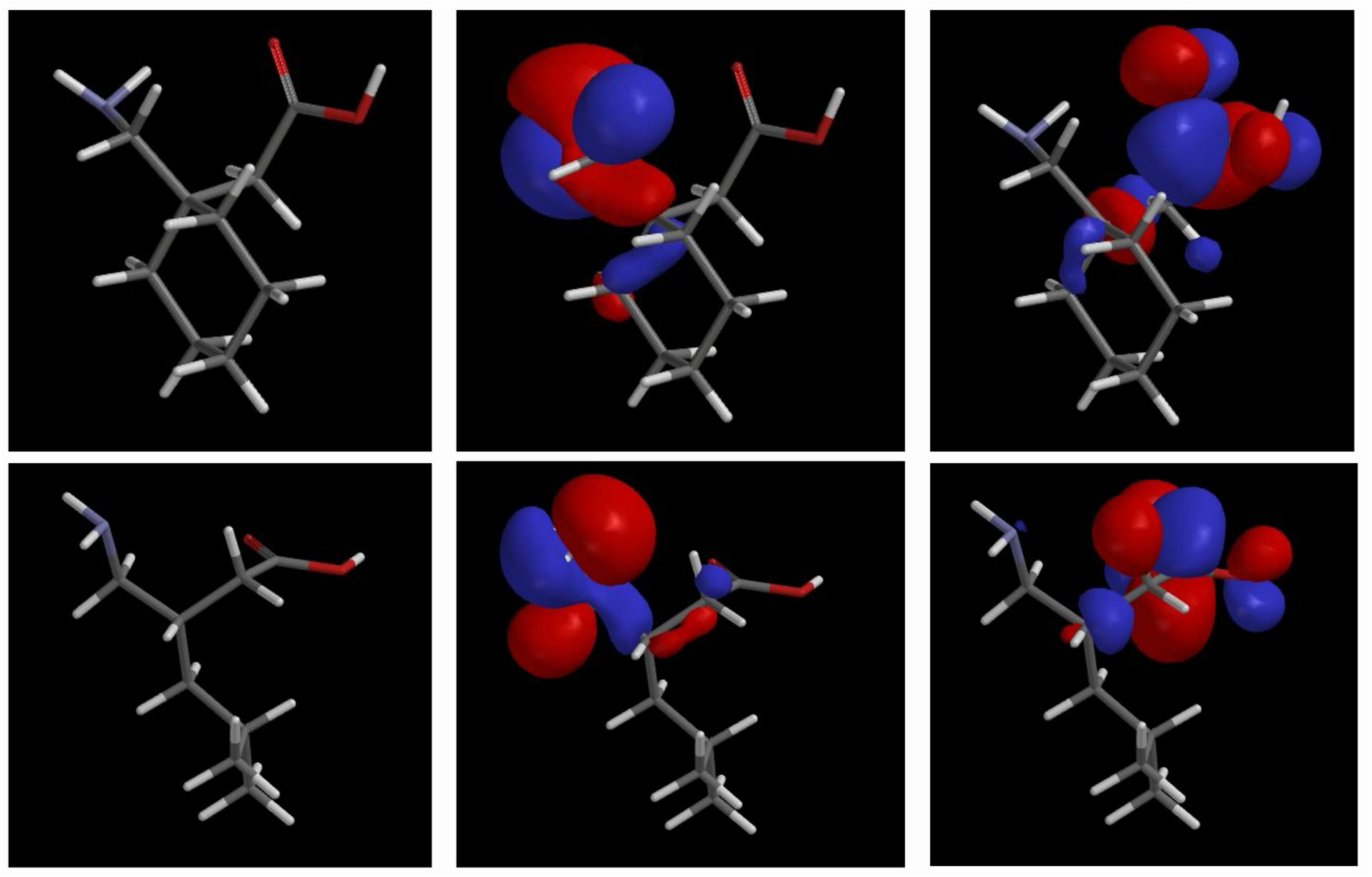
HOMO and LUMO orbital plots of Gabapentin and Pregabalin using DFT method, B3LYP model and 6-31G* basis set, in vacuum for equilibrium geometry at ground state. Molecular orbitals provide important indications about chemical reactivity. The orbital drawing identifies regions where the orbital takes on a significant value, either positive (blue) or negative (red).

## Conclusion

Together, these findings establish that GBP and PGB produce measurable, dose- and time-dependent antinociception in *C. elegans* that is not attributable to experimental confounds in mobility or quadrant preference. Pairing behavioral readouts with direct internal concentration measurements revealed a PK/PD dissociation: at 500 µM, sustained internal drug levels coincided with recovering avoidance behavior, indicating that the late-phase reversal reflects active counter-regulation rather than simple clearance, consistent with our previously reported transcriptional and proteomic signatures. This decoupling parallel documented exposure–response patterns for both gabapentinoids in mammalian systems, supporting a conserved pharmacodynamic feature rather than a species-specific artifact. Molecular modeling further indicates that GBP and PGB share a common electronic pharmacophore for α2δ engagement, while differing in shape descriptors that may plausibly, though not yet mechanistically confirmed, contribute to their divergent absorption and elimination kinetics.

## Acknowledgements

The Université de Montréal partially provided financial support to J. Sultana. A Ph.D. scholarship was awarded to J. Sultana from the *Fonds de recherche du Québec – Santé (FRQS)* https://doi.org/10.69777/347111

## Funding

This project was funded by the National Sciences and Engineering Research Council of Canada (F. Beaudry discovery grant no. RGPIN-2020-05228). Laboratory equipment was funded by the Canadian Foundation for Innovation (CFI) and the Fonds de Recherche du Québec (FRQ), the Government of Quebec (F. Beaudry CFI John R. Evans Leaders grant no. 36706 and 42043). F. Beaudry is the holder of the Canada Research Chair in metrology of bioactive molecules and target discovery (grant no. CRC-2021-00160). This research was undertaken, partly, thanks to funding from the Canada Research Chairs Program.

## Author’s contributions

JS, JDC, JREdC and FB conceived and de-signed research. JS, JDC, JREdC and FB conducted experiments and analyzed data. JS and FB wrote the manuscript. All Authors read, reviewed and approved the manuscript.

## Conflicts of interest

The authors declare no conflict of interest.

## Data Availability

The data generated and analyzed during this study are available from the corresponding author upon reasonable request.

## Supplementary Figures

**Figure S1.**
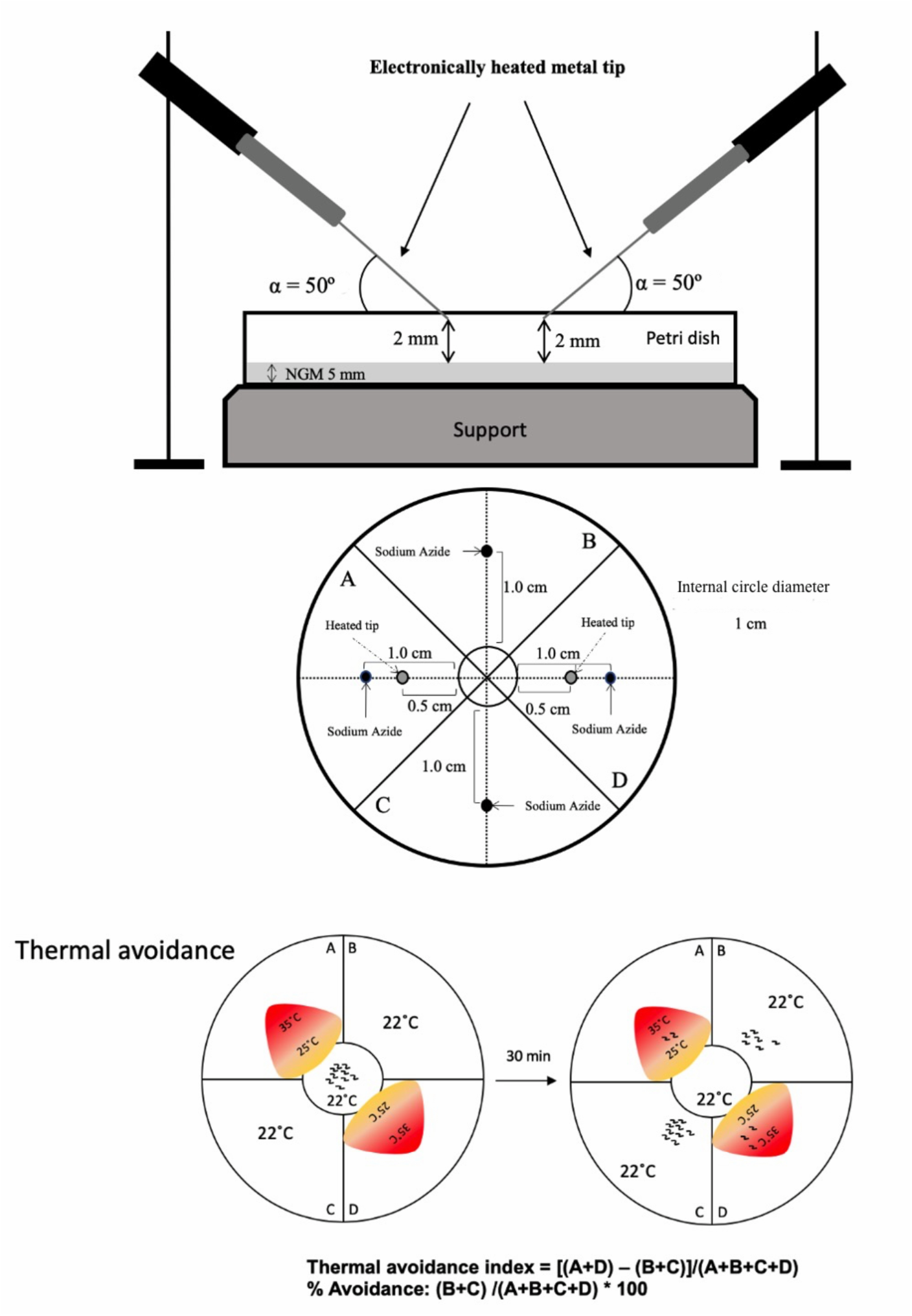
A schematic of the quadrants assay adapted from Margie *et al*. (2013). For head avoidance assay, plates were divided into quadrants two test (A and D) and two controls (B and C). Sodium azide was added to all four quadrants to paralyze nematodes. *C. elegans* were added at the center of the plate (typically, n = 100 to 300) and after 30 minutes, animals were counted on each quadrant. Only animals outside the inner circle were scored. The calculation of thermal avoidance index was performed has described.

**Figure S2.**
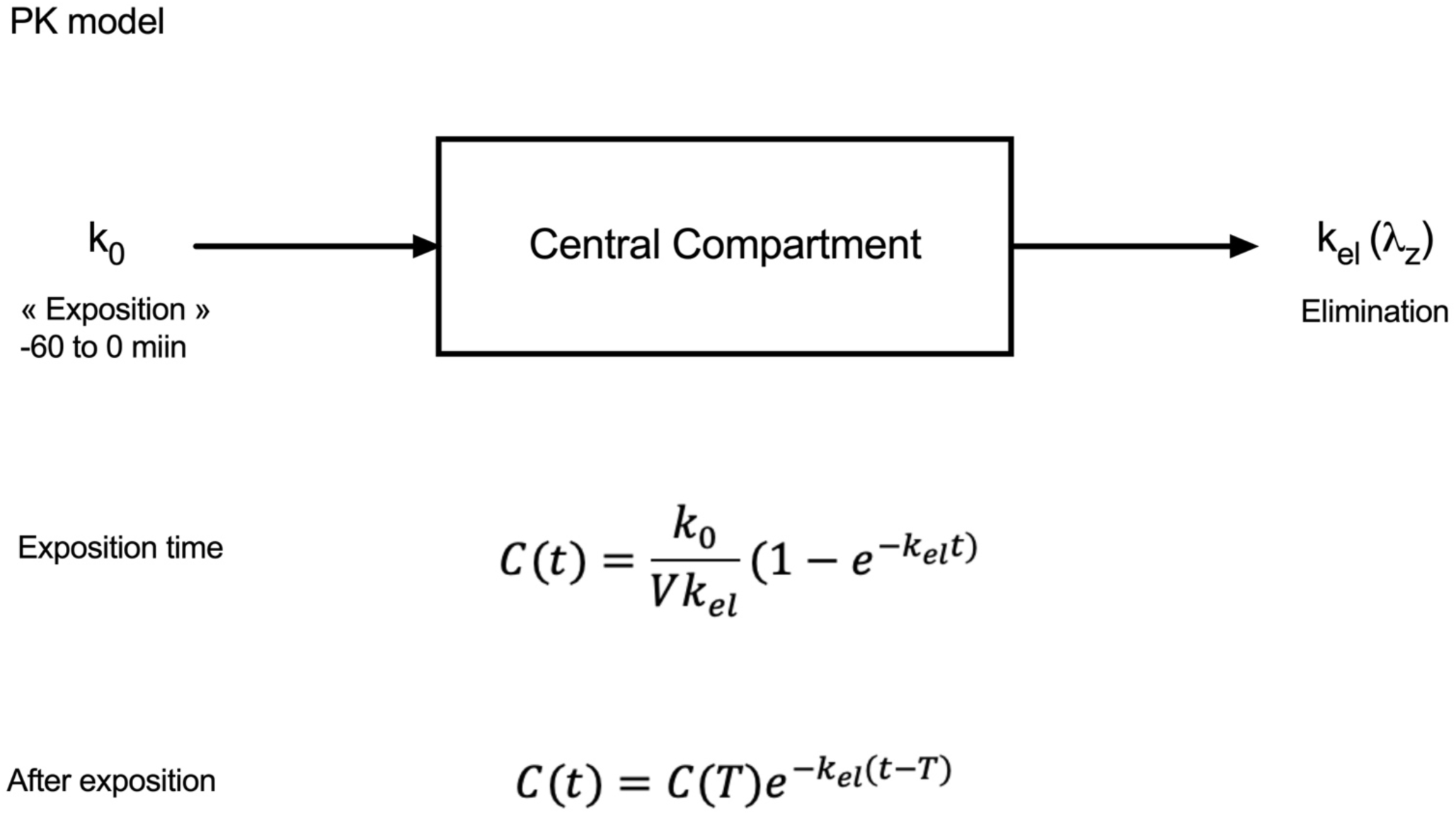
Pharmacokinetic model used to fit gabapentin and pregabalin concentration-time profiles following 1-h exposure of *C. elegans* to 100 µM and 500 µM solutions.

## Notes

### Competing Interest Statement

The authors have declared no competing interest.

